# What The Brain Does As We Speak

**DOI:** 10.1101/2021.02.05.429841

**Authors:** KJ Forseth, X Pitkow, S Fischer-Baum, N Tandon

**Affiliations:** Vivian L. Smith Department of Neurosurgery, McGovern Medical School, Houston, TX, USA; Texas Institute for Restorative Neurotechnologies, University of Texas Health Science Center at Houston, Houston, TX, USA; Department of Neuroscience, Baylor College of Medicine, Houston, TX, USA; Department of Psychological Sciences, Rice University, Houston, TX, USA; Memorial Hermann Hospital, Texas Medical Center, Houston, TX, USA

## Abstract

Language is a defining human behavior and is dependent on networks interactions amongst broadly distributed neuronal substrates. Transient dynamics between language regions that underpin speech production have long been postulated, yet have proven challenging to evaluate empirically. We used direct intracranial recordings during single word production to create a finely resolved spatiotemporal atlas (134 patients, 25810 electrodes, 40278 words) of the entire language-dominant cortex and used this to derive single-trial state-space sequences of network motifs. We derived 5 discrete neural states during the production of each word, distinguished by unique patterns of distributed cortical interaction. This interactive model was significantly better than a model of the same design but lacking interactions between regions in explaining observed activity. Our results eschew strict functional attribution to localized cortical populations, supporting instead the idea that cognitive processes are better explained by distributed metastable network states.

## Introduction

Speech production is a defining human faculty that enables eloquent communication. The ubiquity of word generation with remarkable speed, precision, and fluency belies its complexity. Articulating even a single word requires the selection of a conceptual representation, the construction of a word form, and the orchestration of a complex articulatory plan with associated output monitoring. Despite general agreement on the enumeration of cognitive processes that lead from intention to articulation^1^, no consensus has yet emerged describing the neurobiological architecture that implements these processes.

The cortical basis of speech production has been probed by analyses of deficits secondary to lesions^2^ and neurodegenerative diseases^3^, as well as by analyses of intact language systems through functional imaging^4,5^, structural mapping^6^, and noninvasive electrophysiology^7^. Much of this work has focused on localizing specific cognitive processes to discrete neuroanatomic substrates, yet these efforts have yielded competing interpretations – even in landmark regions like Broca’s area^8^. An alternative framework shifts the focus from patterns of isolated activity in separable substrates to patterns of dynamic interaction between such cortical nodes^9–11^. Evaluation of this theory has been hampered by limitations inherent to the predominant methodologies available for studying the neurobiology of language. These methods lack the temporal or spatial resolution to discern the neural mechanisms driving networks characterized by rapid, transient dynamics across distributed substrates. Invasive human electrocorticography uniquely affords direct, high-resolution recordings of human cortical activity; however, such opportunities are rare and, as a result, prior language studies have been limited in scale and cortical coverage^12–14^.

We overcame these limitations by using large-scale human electrocorticography (134 patients, 25810 electrodes, 40278 trials) to elucidate the neurobiology of language production. This cohort included both subdural surface grid and stereotactic depth electrodes, encompassing the entirety of language-dominant cortex (Figure 1). With this global perspective, we generated a comprehensive spatiotemporal atlas of a classical language paradigm: picture naming with scrambled images as a low-level control. We characterized functionally distinct regions within this atlas by pre- and post-articulatory encoding of established psycholinguistic variables including visual, semantic, lexical, and phonologic correlates. We further developed a grouped dynamical model to resolve discrete neural states that were distinguished by unique patterns of distributed cortical interaction. These data reveal the network architecture of speech production, informing and constraining the neurobiological instantiation of extant language models.

**Figure 1:**
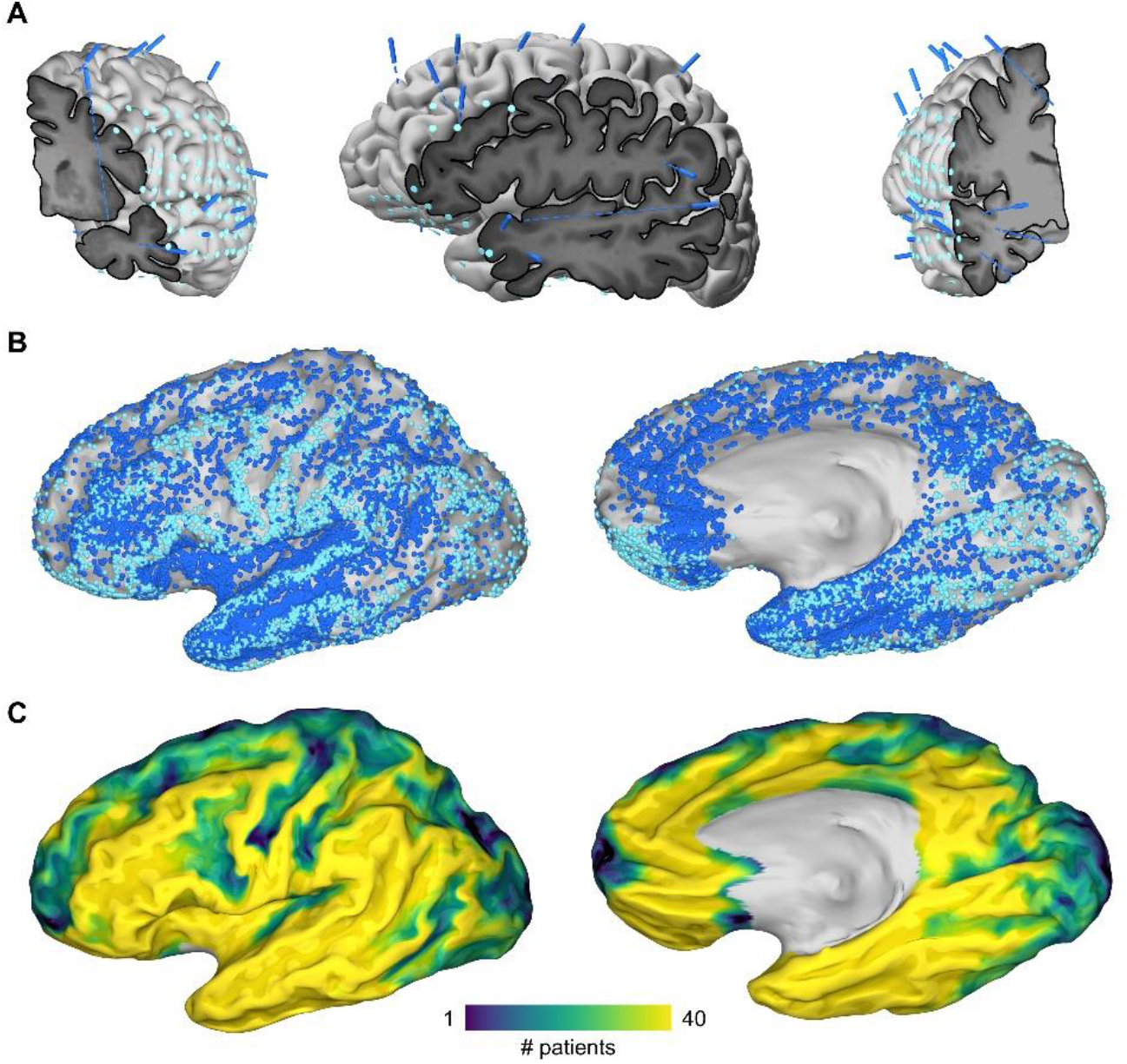
Invasive Human Electrophysiology with Grid and Depth Electrodes in 135 Patients. Distribution of electrodes in left language-dominant cortex. **(A)** Cortical surface model for a single patient who underwent staged implants with grid (light blue) and depth (dark blue) electrodes. The surface has been sliced to reveal intracortical MRI along depth trajectories: frontal-insular (left), supratemporal (middle), medial frontal and midtemporal (right). **(B)** Joint coverage with grid and depth electrodes. Grid electrodes sample from limited cortex at the apex of gyri on the lateral and frontal surfaces; depth electrodes sample evenly throughout cortex including sulci, deep intrasylvian structures, and the medial surface. **(C)** Aggregate cortical coverage in the patient population (n = 134, 25810 electrodes). The surface recording zone for each electrode is estimated by its aspect and location with an inverse model using Euclidean exponential decay subjected to geodesic constraints. 98% of left language dominant cortex is sampled in at least 3 patients.

In addition to providing new insights for language production theory, our approach investigates the broad utility of linking cognitive processes to network states rather than to activity in isolated substrates^15^. We critically evaluate the thesis that complex behaviors comprise sequences of network states^16^, each defining a set of reference dynamics to coordinate the generation and transmission of information throughout the cortex^17^. Speech production is an ideal testbed, requiring the coordination of multiple cognitive systems and resulting in an observable behavior. Elaborating the structural and functional properties of states during speech production will help to guide the derivation of generalizable dynamical principles governing cognition^18^.

## Results

### Global Mean Power Dynamics

Cortical activity was integrated across the cohort to produce a spatiotemporal atlas of cued word production in the language-dominant hemisphere (Figure 2). These global mean dynamics were resolved with surface-based mixed-effects multilevel analysis (SB-MEMA) of high-gamma power in narrow time windows, generating a series of effect sizes and confidence estimates for every point on the standard atlas pial surface. The resulting frames together constitute a high-resolution movie (Figure 2A, Supplementary Figure 1). This analysis was repeated in the language-nondominant hemisphere (Supplementary Figure 2). We further investigated the response properties of this distributed cortical network for naming within regions of interest constrained by the SB-MEMA atlas (Figure 2B). The mean response of each region was analyzed in 3 adjacent time windows: post-stimulus, pre-articulation, and post-articulation (Figure 2C). A broad swath of language-dominant cortex was sequentially recruited during picture naming. The cortical response began in the calcarine sulcus, then spread both dorsally to the intraparietal sulcus and ventrally to the middle fusiform gyrus. Next, a complement of distinct foci in frontal cortex were engaged: pars triangularis, pars opercularis, the supplementary motor area, and the superior frontal sulcus. Significant peri-articulatory activity followed in the subcentral gyrus, peaking with the onset of articulation. The superior temporal and posterior middle temporal gyri were notably quiescent in the pre-articulatory interval, but were engaged throughout articulation.

**Figure 2:**
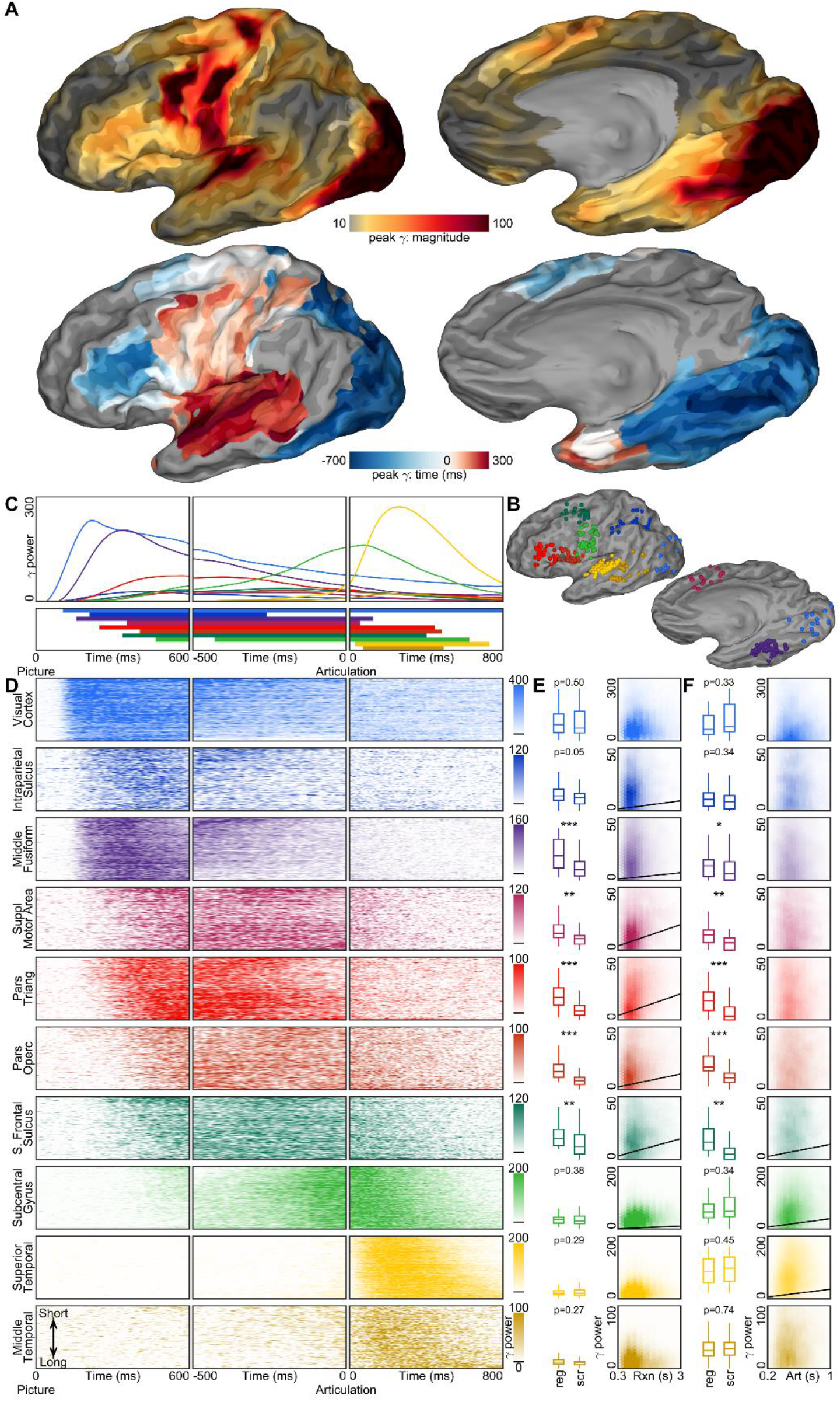
Spatiotemporal Atlas of High-Gamma Power During Picture Naming. Complete mapping of high-gamma power magnitude, extent, and timing in the language-dominant hemisphere. **(A)** Surface-based mixed-effects multilevel analysis of high-gamma power (112 patients, 13812 electrodes, 19465 trials). The model was calculated for 150ms wide windows in 10ms increments from 700ms before to 300 ms after the onset of articulation. For every point on the cortex, the magnitude and timing of peak high-gamma power were identified. The brightness mask highlights cortex that was significantly more active for coherent images than scrambled images during this time window. **(B)** 10 regions of interest were delineated, spanning the picture naming cortical network: occipital (n = 30), intraparietal sulcus (n = 27), middle fusiform (n = 55), supplementary motor area (n = 20), pars triangularis (n = 40), pars opercularis (n = 30), superior frontal sulcus (n = 27), subcentral (n = 41), superior temporal (n = 61), and middle temporal (n = 21). **(C)** The mean percent change in high-gamma power from pre-stimulus baseline in these ROIs is shown for three periods: post-stimulus, pre-articulation, and post-articulation. Significance was determined with the Wilcoxon signed-rank test at an alpha level of p < 0.001 using familywise error correction and further constrained to effect sizes greater than 25% of the peak response. **(D)** Raster plots of single-trial high-gamma power sorted by reaction time in the post-stimulus and pre-articulation windows and by articulatory duration in the post-articulation window. **(E)** Boxplots show average gamma power in the 4 seconds after picture presentation for coherent (left) and scrambled (right) stimuli. Significance was determined with the Wilcoxon rank sum test (^*^ p < 0.05, ^**^ p < 0.01, ^***^ p < 0.001). Hexagonally discretized scatter plots show the distribution of average high-gamma power against reaction time for coherent images. Regression lines are overlaid for those regions with a correlation that was both significant during coherent trials and significantly greater than during scrambled trials. Correlations were evaluated with Spearman’s rho at an alpha level of p < 0.001 and the difference between correlations with Fischer’s z-transform at an alpha level of p < 0.001. **(F)** This analysis was repeated for the average gamma power in the 1 second after articulation onset.

We then quantified the relationship between behavior – reaction time and articulatory duration – and neural response in each region of interest for both coherent and scrambled images (Figure 2D-F). Visual cortex, the intraparietal sulcus, and the middle fusiform gyrus responded at a fixed delay from picture presentation; in contrast, frontal cortex broadly responded later during trials with longer reaction times. Of the regions that were significantly more responsive to coherent images than scrambled images, cumulative high-gamma power and reaction time were significantly correlated in the middle-fusiform gyrus (r = 0.082, p < 10^−13^), pars triangularis (r = 0.278, p < 10^−104^), pars opercularis (r = 0.166, p < 10^−28^), the supplementary motor area (r = 0.082, p < 10^−53^), and the superior frontal sulcus (r = 0.219, p < 10^−42^). Cumulative post-articulatory high-gamma power and articulatory duration were significantly correlated in the superior frontal sulcus (r = 0.072, p < 10^−4^), the subcentral gyrus (r = 0.199, p < 10^−55^), and the superior temporal gyrus (r = 0.182, p < 10^−67^). The middle fusiform gyrus, pars triangularis and opercularis, and the supplementary motor area remained significantly more engaged for coherent versus scrambled images in the post-articulatory timeframe despite having no association with articulatory duration.

This holistic view of the cortical dynamics of picture naming was uniquely afforded by large-scale intracranial electrophysiology encompassing the entirety of language-dominant cortex. While there was a clear progression of activity across the cortex, many regions were jointly active for extended periods. These dynamics are inconsistent with the narrow assignment of linguistic operations to focal and isolated substrates. Instead, they may be better explained by cognitive computation that is orchestrated across transient, distributed, and overlapping networks – a thesis we further evaluate in subsequent analyses.

### Models of Behavior and Neural Response by Linguistic Features

Having characterized the mean spatiotemporal extent of high-gamma power, we evaluated the trial-by-trial effects of distinct linguistic representations on regional activity (Figure 3). Linear mixed-effects models were constructed as a function of visual complexity^19^, semantic familiarity^20^, lexical frequency^21^, lexical selectivity^20^, and phonological density^22^. These models were validated on behavioral measures (Figure 3A): reaction time was best explained by lexical selectivity (ß = 63.54ms, p < 10^−75^), while articulation length was best explained by phonological density (ß = −62.66ms, p < 10^−205^).

**Figure 3:**
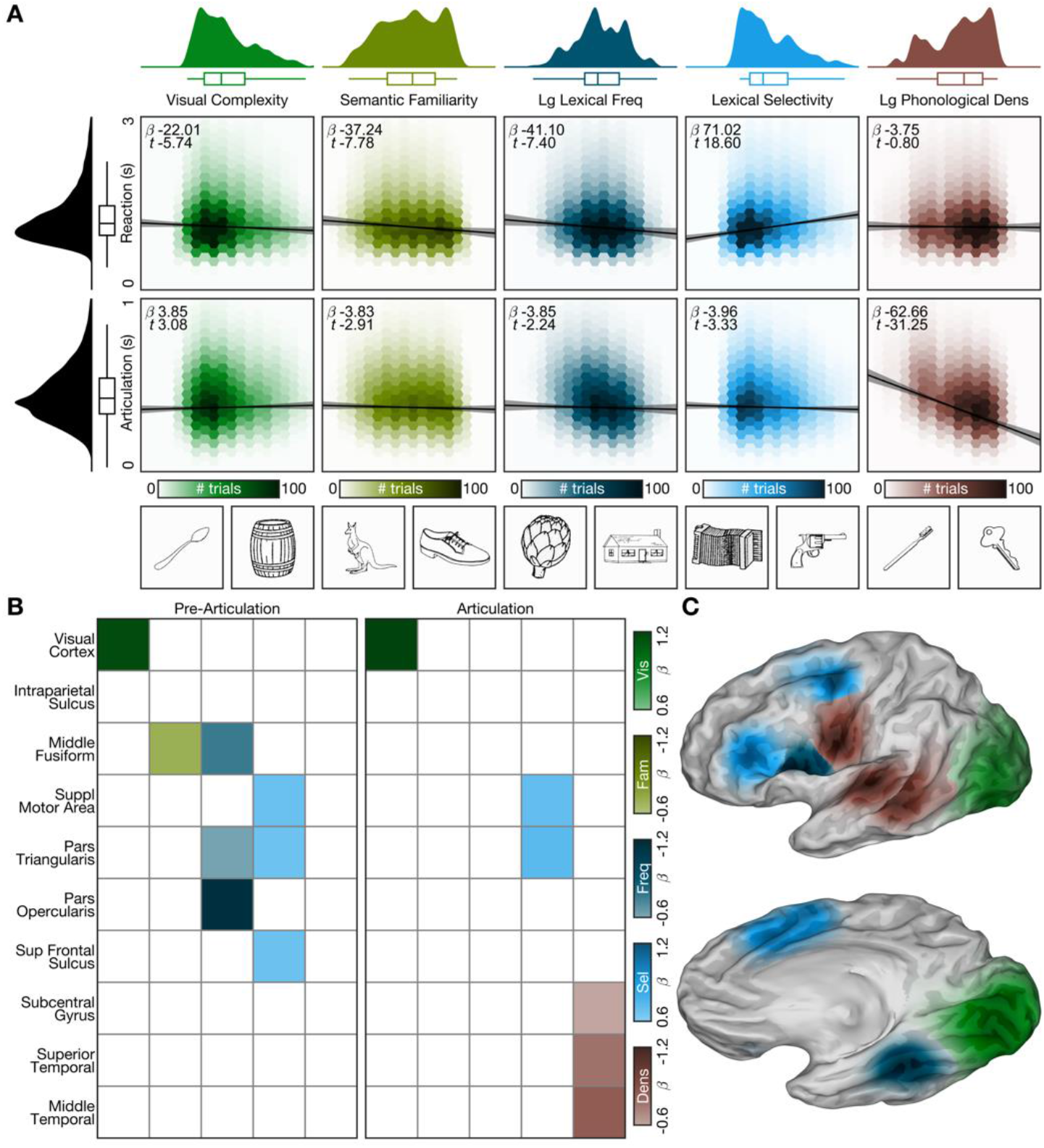
Linguistic Features Mapped to Neurobiology. Linear mixed-effects models of behavior and regional high-gamma power with psycholinguistic predictors. **(A)** Five psycholinguistic parameters were used: visual complexity (SIFT feature count), semantic familiarity (Snodgrass and Vanderwart), log lexical frequency (SUBTLEXus CD count), lexical selectivity (naming agreement), and log phonological density (Irvine Phonotactic Online Dictionary). Reaction time was defined as the duration from picture presentation to the onset of articulation; articulation length spanned from the onset to the offset of articulation. The distribution of each input to the model is shown on the margins. Hexagonally discretized scatter plots include correct trials from patients with electrodes in left language-dominant cortex (112 patients, 19465 trials). The conditional estimates are overlaid with 99% confidence intervals. **(B)** The same parameters were used to model regional high-gamma power response in a pre-articulatory (−2150 to 0ms) and articulatory (0 to 850ms) time window. These ranges were defined as the mean population reaction time and articulation width, respectively, plus twice the standard error of the mean. Significance was evaluated at an alpha level of p < 0.01. **(C)** The aggregate cortical spread of electrodes in each region of interest is colored by the predictor with the largest significant *t*-value.

We applied these models to high-gamma power in each region of interest during time windows before and after the onset of articulation. Visual cortex activity was related to visual complexity of the stimulus (ß = 0.116, p < 10^−5^), consistent with localized feature processing. In the pre-articulatory period, semantic familiarity was uniquely related to middle fusiform gyrus activity (ß = −0.071, p < 10^−3^). Lexical frequency was also encoded in the middle fusiform gyrus (ß = −0.077, p < 10^−4^), as well as in pars triangularis (ß = −0.063, p = 0.0053) and opercularis (ß = −0.116, p < 10^−4^). Lexical selectivity was related to activity in pars triangularis (ß = 0.069, p < 10^−3^), the supplementary motor area (ß = 0.070, p = 0.0014), and the superior frontal sulcus (ß = 0.069, p = 0.0011). After the onset of articulation, phonologic density of the spoken response was encoded in perisylvian regions: subcentral (ß = −0.061, p < 10^−4^), superior temporal (ß = −0.078, p < 10^−8^), and posterior middle temporal gyri (ß = −0.085, p = 0.0043).

### Interregional Interactions of Cortical State Dynamics

We have established that language engages a distributed network of regions with concurrent activity and separable linguistic correlates; however, our analyses thus far assume that cognitive operations are locally computed in isolated substrates and time-locked to observable events. This assumption is deeply embedded in the cognitive neuroscience literature, confounding attempts to distinguish the neurobiological correlates of interactivity that are foundational to psycholinguistic models of speech production (Supplementary Figure 3)^23^. To relax this assumption and directly resolve interactions between regions, we implemented a grouped autoregressive hidden Markov model (gARHMM, Figure 4A). Critically, this model learns a single set of latent dynamical parameters across the patient population and emits state sequences of network interactions for each trial^24^.

**Figure 4:**
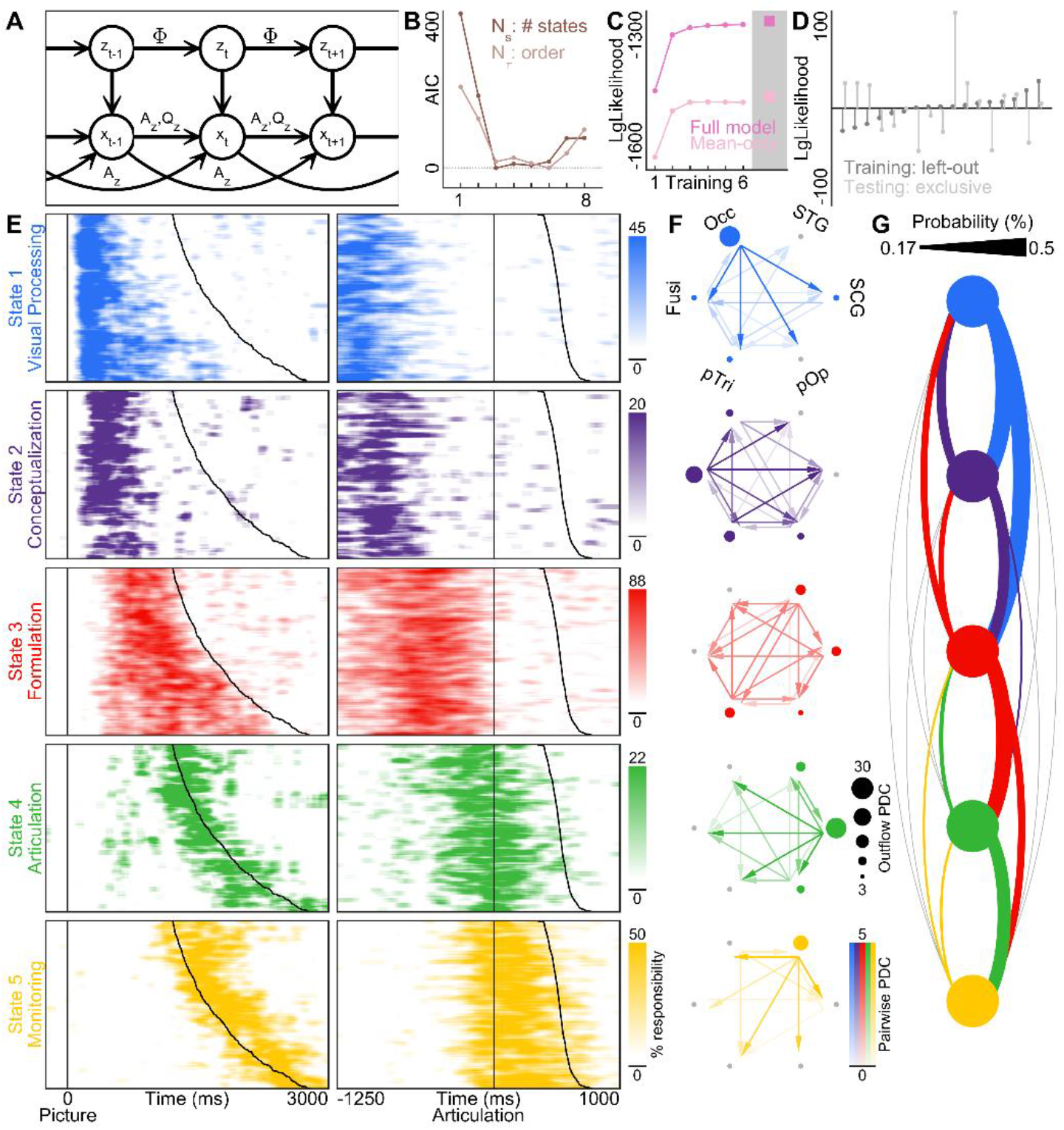
Nonlinear Dynamics in Distributed Cortical Networks. Autoregressive hidden Markov model applied to single-trial high-gamma power in 17 patients with electrodes over visual cortex, middle fusiform gyrus, pars triangularis and opercularis, subcentral gyrus, and superior temporal gyrus. **(A)** Model design schematic. Latent states *z* and observations *x* evolve according to autoregressive state dynamics *A*, regional process covariances *Q*, and state transition probabilities Φ. A single set of parameters *A, Q*, and Φ were learned across all 19 patients. **(B)** Hyperparameter selection with 10-fold cross-validation on the training set, evaluated with AIC relative to trough. **(C)** Model comparison. Training and cross-validated testing for fully interactive model and reduced mean-only model. **(D)** Analysis of robustness and validity. Each patient was iteratively held-out from training. The full model was then tested on trials exclusively from each patient. The model is robust to any single patient and generalizes well to all patients. **(E)** State sequences on held-out testing data. Each of the 5 active states is shown with a unique color. For each raster, all trials were sorted by reaction time in the left column and by articulatory duration in the right column. The maximum of each state color scale is set to 400% of the mean state density. **(F)** Interregional dynamics. The graph nodes represent distinct regions. The graph edges represent the pairwise partial directed coherence (PDC), a Granger causal measure of interaction derived from the unique dynamics that define each state. The size of each node is proportional to the total outflowing PDC. The significance of every pairwise interactional coefficient and each nodal outflow was evaluated via bootstrapping (n = 10000) at an alpha level of p < 0.001. No interactional coefficient in the residual state was significant. **(G)** State transition probabilities. The observed probability of transitioning “forwards” (e.g. blue to purple) is shown flowing downwards on the right; probability of transitioning “backwards” (e.g. yellow to green) is shown flowing upwards on the left. The lower threshold is set to the uniform probability of non-self-transitions.

17 patients had concurrent coverage of the core language network: visual cortex, middle fusiform gyrus, pars triangularis, pars opercularis, subcentral sensorimotor cortex, and superior temporal gyrus. The hyperparameters – model order (τ = 3) and number of states (*N_z_* = 6) – were determined by 10-fold cross-validation (Figure 4B) of the training dataset (80% of trials uniformly sampled from all patients). Latent dynamical parameters were then optimized on the training dataset and applied to the held-out test dataset. This model performed significantly better (p < 10^−162^) than a model of the same design but lacking interactions between regions (Figure 4C). Model training was unbiased by data from any single patient and testing generalized to data from held-out patients (Figure 4D).

We identified 6 dynamical states – 5 states during speech production and a background state between trials (Figure 4E). The 5 active states demonstrated a consistent temporal precession relative to both picture presentation and articulation. These states lend themselves in number and timing to established psycholinguistic nomenclature^1^: visual processing, conceptualization (activation and selection of a lexical concept), formulation (staged form encoding to produce gestural scores), articulation, and monitoring (Supplementary Figure 4). The Granger-causal interactions between regions that define the dynamics of each state were quantified with partial directed coherence (Figure 4F). A second model trained on trials with scrambled images also identified 5 active states, but with differences in network architecture and state frequency (Figure 5, Supplementary Figure 5).

**Figure 5:**
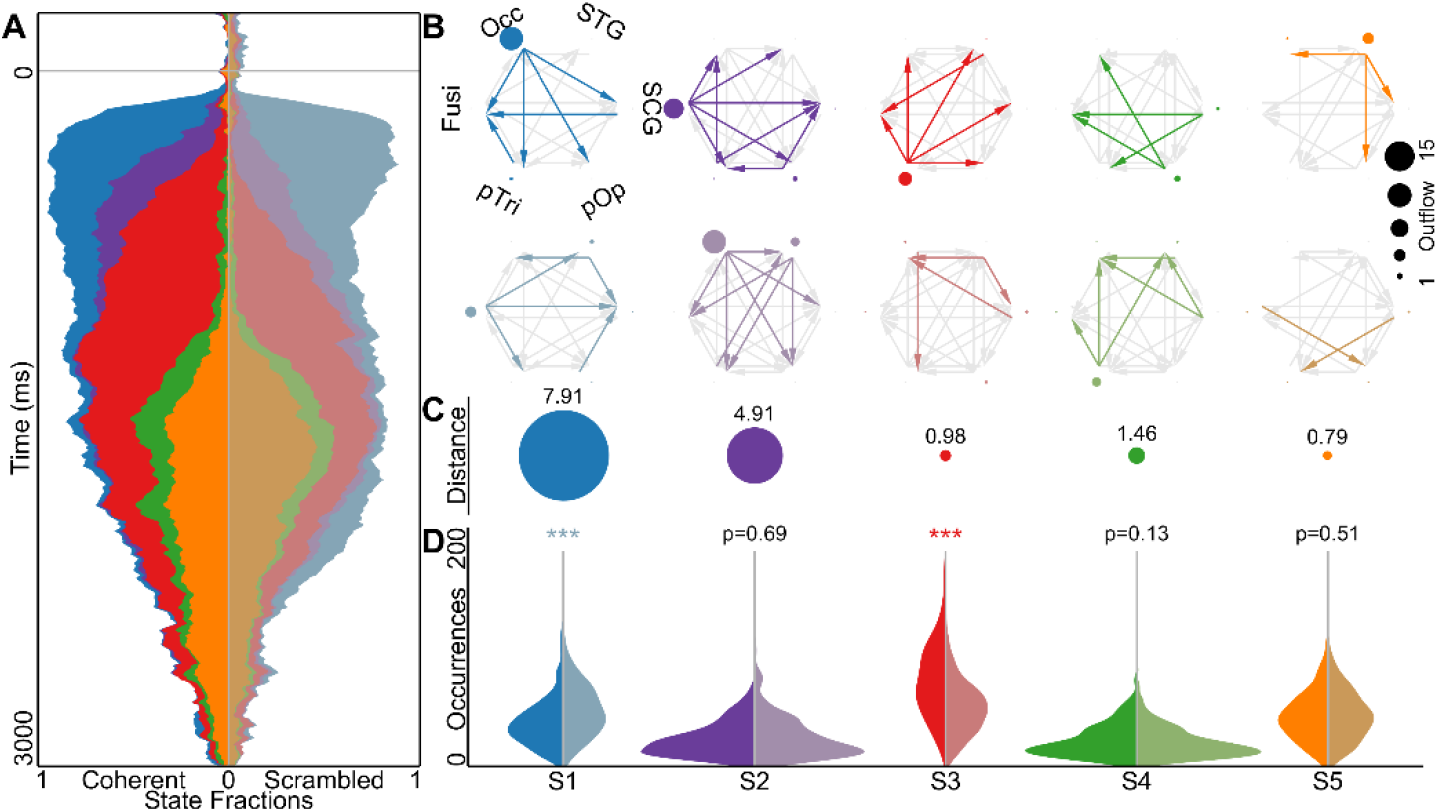
Network Activity Contrast in Coherent and Scrambled Picture Naming. Distinct nonlinear dynamics in cortical networks during trials with coherent versus scrambled pictures. **(A)** The fraction of trials engaging a given state as a function of time from stimulus presentation for coherent (left, light colors) and scrambled (right, muted colors) images. **(B)** Contrast of the learned network states. Pairwise interactions, quantified with partial directed coherence, that were greater during coherent trials are shown in the top row; those greater during scrambled trials are shown in the bottom row. Similarly, nodes with greater outflow during coherent versus scrambled trials are shown in the top and bottom row, respectively, with radius proportional to the magnitude of difference. The significance of every pairwise interactional coefficient and each nodal outflow was evaluated with the Wilcoxon rank sum test on bootstrapped distributions (n = 10000) at an alpha level of p < 0.001. **(C)** Cosine similarity of each network state pair. The first two states were different during coherent and scrambled images, while the latter three states were similar. **(D)** Distribution of state frequency. Significance was evaluated with the Wilcoxon rank sum test (^*^ p < 0.05, ^**^ p < 0.01, ^***^ p < 0.001).

Each state featured significant contributions from a complex network of pairwise regional interactions that were essential for optimal modeling of neuronal dynamics; the most salient interactive properties of these states are described here. The first state, *visual processing*, was concentrated in the ∼250 ms immediately following picture presentation and its dynamics were dominated by outflow from visual cortex. This was followed by the second state, *conceptualization*, from ∼250 to ∼500 ms in a distributed network organized by outflow from the middle fusiform gyrus and pars triangularis. The network architecture of these two states was significantly altered during scrambled trials (Figure 5C); in particular, the outflow from the middle fusiform gyrus was largely replaced by outflow from visual cortex. The third state, *formulation*, recruited a decentralized perisylvian network that remained engaged through the onset of articulation and accounted for the majority of variance in reaction time (ß = 7.41, p < 10^−23^). The network architecture of this state was similar for coherent and scrambled trials, but it occurred significantly more frequently in response to coherent images (p < 10^−23^, Figure 5D). The fourth state, *articulation*, was engaged in the ∼400 ms around articulatory onset with dynamics dominated by outflow from subcentral sensorimotor cortex. The fifth and final state, *monitoring*, was active throughout articulation relied predominantly on outflow from superior temporal gyrus. Both *articulation* and *monitoring* featured convergent network architectures and state frequencies for coherent and scrambled images. These results are consistent with our thesis that the fundamental unit of linguistic computation in the brain is not a set of discrete functional regions, but a sequence of dynamical network states.

We also computed state transition probabilities that describe the rate of observed pairwise switching (Figure 4G). These revealed a locally interactive state switching behavior conserved across trials of varying reaction time and articulatory length. For every state, the most likely transition was to the next state. Transitions from formulation to any other state were common, while transitions directly from visual processing or conceptualization to articulation or monitoring were below chance. Together, the restricted set of cognitive state transitions and the imbricated set of interactive state dynamics ground the discreteness-interactivity axis^23^ of speech production models^1,25,26^ in the neurobiology of human cortex.

## Discussion

### Spatiotemporal Cartography of Language Dominant Cortex

Language involves the coordinated activity and interaction of diverse cortical substrates, yet prior electrocorticographic investigations of language function have studied this broad network in fragments^13,14,27–30^ due to the unique challenges of invasive human electrophysiology that limit the scope and scale of recordings^31–33^. Complete coverage of the cortical surface with intracranial electrodes requires upwards of 10,000 contacts^34^. Furthermore, grid and depth electrodes have distinct stereotypic coverage probabilities: grid electrodes predominantly record from gyri on the lateral or ventral surface, while depth electrodes are more likely to record from cortical sulci, the ventral temporal surface, the medial bank of the hemisphere, and the insula. Our study yielded an unprecedented database of intracranial recordings from both grid and depth electrodes with the magnitude, density, and homogeneity of coverage necessary to build a comprehensive spatiotemporal map of speech production encompassing the entirety of language-dominant cortex. We used this resource to finely characterize the activity at specific network nodes and the evolving interactive dynamics between nodes. Such an approach is imperative for definitively resolving the processes that lead from intention to articulation^1^.

### From Intention to Articulation

Despite general agreement on the processes required to convert a picture to its spoken name^26,35^, the instantiation of these processes in neural dynamics is unknown. Psycholinguists and neurobiologists have long engaged in parallel approaches for the study of speech production, probing the hidden internal processes of language generation and exploring the functional contributions of discrete neuroanatomic substrates. Picture naming has demonstrated exceptional utility in these pursuits. Varying stimulus complexity of isolated representational levels – semantic, lexical, and phonological – yields behavioral data which outline the veiled architecture of speech production. Reaction time variations following challenges to the production system with level-specific interference at early and late windows suggest at least two separate steps in lexical access^25,36,37^. Error types and rates constrain the interactions between representational levels in formalized models of production^38–40^ and, through the anatomic correlates of distinct aphasic patterns, establish connections to neurobiology^41^. Functional imaging in subjects with intact language, derived from measures of blood flow and metabolism, has catalyzed the spatial categorization of nodes in a complex and distributed language network^5^. The breadth of recruited substrates argues against localizationist accounts of production^42,43^ in favor of network-dependent cognitive processes^44,45^, but measuring internodal communication requires fine temporal scale. Electrophysiology directly captures the dynamical behavior of neural substrates with radically improved temporal resolution^46–48^. In compiling a large dataset of intracranial recordings during picture naming, we are able to integrate cognitive models of speech production with the neurobiology of distributed and interactive cortical networks. We reveal 5 states engaged during picture naming, remarkably concordant with the 5 cognitive stages named in the seminal work by Indefrey and Levelt: visualization, conceptualization, formulation, articulation, and monitoring^1^. Each of these states is dually characterized by timing within a trial-specific transition sequence and by functional network structure comprising directed information exchange.

The state characteristics inform two fundamental and disputed properties of conflicting production models: the seriality of separable cognitive processes and the interactivity between representational levels^23,26,49^. Our findings are concordant with a concerted serial propagation of network dynamics. A consistent sequence of states is observed from picture presentation through articulation and state switching is tightly constrained within pre- and post-formulation periods. Distant interactions, reflecting the unique pattern of information exchange during each state, underline the distributed basis of cognitive processes. Some states, such as visualization and monitoring, are directed by singular foci while others feature a balanced distribution of largely reciprocal connections, as epitomized by formulation. Local interactions are manifest in substrates shared between states – most notably in ventral lateral prefrontal cortex during conceptualization, formulation, and articulation. These reflect a functionally heterogeneous population of neurons contributing to the signal measured at each electrode. Our results comprehensively delineate the neurobiology of picture naming in language-dominant cortex, robustly motivate the use of seriality in computational models of speech production, and establish a concrete mechanism for representational interactions in language networks.

### States Approximate Nonlinear Dynamics

We present two complementary accounts of temporal dynamics. The first, mean gamma activity, is local and rooted in physical space; the second, network connectivity patterns, is global and defined in state space. In a similar manner to piecewise linear approximation of a curve, this second account uses the switching characteristic of the hidden Markov model to approximate the high-dimensional state space of neural dynamics^50,51^. Each state then represents a set of reference dynamics at informative inflections of state space. Fluctuations around these reference dynamics provide a generic mechanism by which to disseminate information in a structured manner throughout complex networks^17^. The pairwise measures of information flow that we quantify are thus an average of interregional exchanges, amalgamating transmissions between small groups of neurons. This perspective integrates distributed interactions^52^ with modular cognitive processes^1^.

Neural state sequences comprise a powerful framework by which to model cognition. We provide empirical evidence consistent with this framework; our dynamical model identifies 5 states in speech production. Specifying fewer states results in refolding a pair of states together; specifying more states results in degenerate splitting of the baseline state. Stated another way, simplifications of state space produce coarser structures that exhibit functionally similar behavior^53^. A hierarchically organized neural state structure may reveal increasing dynamical detail with improved observation of the system. This behavior suggests that incorporating data from additional regions, latent parameters for patient-specific network variability, and progressively finer-scale cortical recordings could result in further decomposition of our observed states, e.g. formulation might splinter into morphologic, phonologic, and phonetic encoding. Uncovering dynamical systems in the brain provides an improved understanding both of granular processes, such as picture naming, and of cognition more generally.

## Materials and Methods

### Experimental Design

We recorded cortical activity in 134 patients undergoing presurgical evaluation of refractory epilepsy with either subdural grid electrodes (46 patients, 6120 electrodes) or stereotactic depth electrodes (99 patients, 19690 electrodes). Patients named images of common objects; a scrambled image control condition was interweaved. Stimulus, naming accuracy, articulation onset and offset, and articulatory content were recorded for every trial. A surface-based mixed-effects multilevel analysis (SB-MEMA) of high-gamma activity was used to generate a high-resolution whole-brain movie of mean power dynamics. To characterize the function of local responses in nodes of the distributed naming network, the regional variance in high-gamma power during pre- and intra-articulatory time windows was explored with canonical psycholinguistic descriptors. Single-trial nonlinear interactional dynamics of ventral and perisylvian cortex were subsequently described with the novel application of a grouped autoregressive hidden Markov model (gARHMM). Model hyperparameters were optimized by cross-validation on held-out data.

### Human Subjects

After obtaining written informed consent, we enrolled 134 patients (65 men, 69 women; mean age 33 ± 10 years; mean IQ 96 ± 14) undergoing evaluation of intractable epilepsy with intracranial electrodes. The study design was approved by the committee for the protection of human subjects at The University of Texas Health Science Center. Hemispheric language dominance was evaluated in 88 patients (left, n = 82; right, n = 6) by intracarotid sodium amytal injection^54^ (n = 40), fMRI laterality index^55^ (n = 27), or cortical stimulation mapping^56^ (n = 21). The remaining patients were determined to be right (n = 43) or left handed (n = 3) by the Edinburgh Handedness Inventory^57^; they were assumed to be left hemisphere language-dominant.

### Language Paradigm

Patients engaged in a picture naming task. They were instructed to articulate the name for common objects depicted by line drawings^20,58^ as quickly and accurately as possible. A control condition was intermixed consisting of the same images with pixel blocks randomly rotated; patients were instructed to respond with “scrambled.” Each visual stimulus was displayed on a 15-in LCD screen positioned at eye level for 2 seconds with an interstimulus interval of 3 seconds. A minimum of 120 (mean 298) visual stimuli were presented to each patient using stimulus presentation software (Python v2.7). Mean accuracy was >90% in all patients.

### Structural Imaging

Preoperative anatomical MRI scans were obtained using a 3T whole-body MRI scanner (Philips Medical Systems) fitted with a 16-channel SENSE head coil. Images were collected using a magnetization-prepared 180° radiofrequency pulse and rapid gradient-echo sequence with 1-mm sagittal slices and an in-plane resolution of 0.938 × 0.938 mm. Pial surface reconstructions were computed with FreeSurfer (v5.1)^59^ and imported to AFNI^60^. Postoperative CT scans were registered to preoperative MRI scans for localization of electrodes relative to cortex. Grid electrode locations were determined by a recursive grid partitioning technique and then optimized using intraoperative photographs^61^. Depth electrode locations were informed by implantation trajectories^62^ from the Robotic Surgical Assistant (ROSA, Zimmer-Biomet) system.

### Electrophysiology Acquisition

Grid electrodes—subdural platinum-iridium electrodes embedded in a silicone elastomer sheet (PMT Corporation, top-hat design; 3-mm diameter cortical contact)—were surgically implanted via a craniotomy^56^. Electrocorticography recordings were performed at least 2 days after the craniotomy to allow for recovery from the anesthesia and narcotic medications. Depth stereo-electroencephalographic platinum-iridium electrodes (PMT Corporation; 0.8-mm diameter, 2.0-mm length cylinders; separated from adjacent contacts by 1.5 to 2.43 mm) were implanted using ROSA, with stereotactic skull screws registered to both a computed tomographic angiogram and an anatomical MRI^63–65^. There were from 8 to 16 contacts along each depth probe, and each patient had multiple (12 to 18) probes implanted.

For the first 19 recordings, electrocorticography data were collected with a sampling rate of 1 kHz and bandwidth of 0.15 to 300 Hz using Neurofax (Nihon Kohden). For the latter 126 recordings, data were collected with a sampling rate of 2 kHz and bandwidth of 0.1 to 700 Hz using NeuroPort NSP (Blackrock Microsystems). Continuous audio recordings of each patient were performed with both an omnidirectional microphone (Audio Technica U841A, 30 to 20,000 Hz response, 73 dB SNR) placed adjacent to the presentation laptop and a cardioid lavalier microphone (Audio Technica AT898, 200 to 15,000 Hz response, 63 dB SNR) clipped to clothing near the mouth. These recordings were analyzed offline to transcribe patient responses, as well as to determine the time of articulatory onset and offset.

### Data Processing

A total of 25810 electrodes (6120 grids, 19690 depths) were implanted in this cohort (Figure 1). Only the 18483 electrodes (4476 depths, 14007 grids) unaffected by epileptic activity, artifacts, or electrical noise were used in subsequent analyses. Each trial was defined by the presentation of a visual stimulus. Trials in which the patient answered incorrectly or did not respond were eliminated. Additionally, trials in which the patient responded after more than 3 seconds were removed.

### Digital Signal Processing

Analyses were performed with trials time-locked to either picture presentation or to articulation onset. In all analyses, the baseline was defined relative to the picture presentation (−750 to −250 ms). Line noise was removed with zero-phase second-order Butterworth bandstop filters at 60, 120, and 180 Hz. The high-gamma (60 to 120 Hz) analytic signal was generated from raw electrocorticography data by a frequency domain bandpass Hilbert filter (paired sigmoid flanks with half-width 1.5 Hz)^62,66^. Power was then calculated as the squared envelope of the analytic signal, normalized as a percent of baseline activity.

### Inverse Model of Electrode Recording Zones

In the same manner as previous work^29,62^, response properties of individual electrodes were mapped to patient-specific cortical models via a surface recording zone. This zone both constrained the surface-based spatial registration of individual cortical models to a standard atlas and constrained the weighted estimate of cortical contributions to the observed signal at each electrode.

The definition of electrode recording zone was tailored to the type of electrode. For grid electrodes, the central coordinate of each electrode was matched to its closest node on the cortical envelope. This seed was then grown to a geodesic radius of 1.5 mm, matching the dimensions of the electrode. Each of the vertices within this region was mapped to its closest vertex on the pial surface model. These seeds were then grown along the surface to a maximum geodesic radius of 10 mm, constituting the surface electrode recording zone. For depth electrodes, the central coordinate of each electrode was simply mapped directly to the closest vertex on the pial surface model. This seed was then grown to a maximum geodesic radius of 10 mm to define the surface electrode recording zone.

For both electrode types, the inverse model within the recording zone was defined by the same piecewise inverse function. Cortex directly adjacent to electrodes (e.g. beneath grid electrodes or alongside depth electrodes) was assigned a weight of 1; more distant cortex was weighted according to exponential decay with a full-width half-maximum at 2.3 mm. Individual electrode statistics were subsequently propagated onto the cortical surface with this inverse function.

### Surface-Based Registration to Standard Atlas

All group analysis was performed in standard space on the MNI N27 cortical surface. Electrode locations and recording zones were transformed to standard space with a nonlinear surface-based registration^67,68^. This registration was used to generate coverage maps, define regions of interest, and to enable group statistics at each vertex in the mixed-effects multilevel analysis^29,62^.

### Mixed-Effects Multilevel Analysis

To generate statistically robust and topologically precise estimates of percent change in power across the cortex, we used a surface-based mixed-effects multilevel analysis (SB-MEMA, Supplemental Figure XX) leveraging the electrode recordings zones and nonlinear transform to standard space defined previously. This method addresses universal challenges for grouped analysis of human invasive electrophysiology: a) accurate localization and spatial localization of cortical sources^29,62^; b) data integration across the cohort, accounting for sparse sampling and anatomic variability^67–69^; c) statistical modeling of population-level effects, mitigating outlier inferences and accounting for intra- and interpatient response variability that violate the assumptions of simpler models (e.g., ANOVA, t-tests)^28,70^.

The model consisted of two levels: the individual and the group. At the individual level, an estimate of percent change in power at each electrode was fitted with a mixed-effects model informed by the sampling variance. The resulting effect and significance estimates were propagated onto the patient-specific cortical model using the surface recording zone of each electrode. These patient-specific maps then underwent surface-based registration to the standard atlas. Finally, a mixed-effects model at the group level generated the effect and significance estimates for each vertex on the MNI N27 atlas^29,62^. The entirety of this model is graphically described in Supplemental Figure 6.

This model was run on 150 ms wide windows at 10 ms increments separately aligned to picture presentation and articulation onset. These were then integrated into two continuous movies^48^ – one for each alignment. A geodesic Gaussian smoothing filter (3 mm full-width at half-maximum)^62^ was applied to each frame. Significance was evaluated at an alpha of p < 0.01 with familywise correction for multiple comparisons^29^. Activity masks were further constrained to coverage in at least 3 patients and effect estimates exceeding a 5% change in power^62^.

### Regions of Interest Delineation

To further explore the distributed cortical dynamics of naming, we delineated 10 regions of interest by functional and anatomical constraints. The full SB-MEMA model in the language dominant hemisphere revealed spatiotemporally distinct nodes of activity. These were further refined by anatomical boundaries (e.g. inferior frontal gyrus was split into pars triangularis and opercularis). Each region of interest was explicitly defined by a central point in standard MNI N27 space and a geodesic radius on the atlas pial surface. Electrodes within each region were identified; those that were deemed inactive with a high-gamma power percent change of less than 10% from baseline were removed. For patients with multiple electrodes within a single region of interest, only the most active electrode was retained.

### Grouped Time Series Analysis

Grouped estimates of the percent change in power for a regional collection of electrodes were calculated by averaging across trials at each electrode and then determining an ensemble average across electrodes within each region^28,62^. A Savitsky-Golay polynomial filter (3rd order, 251-ms frame length) was subsequently applied to each time series. Significance was evaluated against baseline and between conditions with two-sided paired t-tests corrected for multiple comparisons with the false discovery rate.

### Selection of Psycholinguistic Features

Four psycholinguistic features were chosen to represent distinct cognitive processes in the production of single-word articulations. Each is well-studied and readily available to optimize cross-study comparisons. Visual complexity was assessed for each stimulus with the number of Scale Invariant Feature Transform descriptors^19^. Familiarity and selectivity were assessed for each stimulus using the Snodgrass and Vanderwart measures^20^. Lexical frequency was assessed for each articulation using the SUBTLEXus CD count^21^. Phonologic density was assessed for each articulation, quantifying the number of words with a unitary Hamming distance from the target^22^. The expected psychometric effects of these variables on reaction time and articulatory duration were validated with separate linear mixed-effects models.

### Regional Linear Mixed Effects Models

For each of the 10 regions of interest, the variability of single-trial high-gamma power was modeled with the psycholinguistic features described earlier. Total gamma power in 4 time windows of interest were separately modeled. Each model was comprised of fixed effects for each psycholinguistic feature, a random intercept grouped by patient, and a random estimate of each psycholinguistic feature grouped by patient. All effect sizes describe the change per standard deviation.

### Grouped Autoregressive Hidden Markov Model

The autoregressive hidden Markov model (ARHMM) combines autoregressive (AR) stochastic linear dynamics with the discrete switching latent states of a hidden Markov model (HMM)^24^. This method enables single-trial analysis that does not require manual alignment of trials by picture presentation or articulation onset. Furthermore, it obviates the dubious assumption underlying all cross-trial averaging metrics that the same cognitive processes are occurring at the same times in all trials. All latent parameters – the time series of network states, their transition probabilities, and the dynamics of each state – are inferred directly from neural data. In this work, we have expanded on our past instantiation of this model with the implementation of a grouped training method (gARHMM) that enables us to infer a single set of generalized latent parameters across the entire patient cohort.

An AR process is a random process with temporal structure, where the current state *x*_*t*_ of a system is the sum of a linear combination of previous states and a stochastic innovation *v*_*t*_drawn from zero-mean isotropic white noise. The linear dynamics of such a system can be described by *N*_τ_ matrices of AR coefficients *A*_τ_ at different time delays τ, which can be combined into a tensor ***A*** = {*A*_τ_}. The stochastic elements of the AR process are specified by a covariance matrix *Q*.

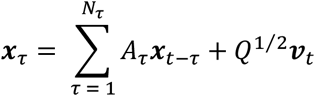

Since this model is linear, it is a poor approximation of nonlinear neural dynamics. This limitation motivated our subsequent extension of the model to include switching dynamics.

The defining property of a first-order Markov model is that transition probabilities between states depends only on the previous state. In a *hidden* Markov model, this dynamic is unobserved – each state emits observable quantities with some associated probability. *Autoregressive* hidden Markov models combine the stochastic linear dynamics of AR processes with the partial observability of a hidden Markov model. Here we use a discrete latent state z and assume autoregressive Gaussian emissions conditioned on that latent state. Each latent state z indexes a different stochastic linear process with a state-specific dynamics tensor ***A**_z_* and a state-specific process noise covariance *Q_z_*. The switching characteristics allow an ARHMM to approximate a stochastic nonlinear dynamical system.

In the context of invasive electrophysiology, the observations are time series of high-gamma power at a fixed number of regions (visual cortex, mid-fusiform gyrus, pars triangularis, pars opercularis, subcentral gyrus, and superior temporal gyrus). The AR coefficients *A_zτjk_* specify the Granger causal dynamical relationship between regions *j* and *k* at time lag τ in state *z*. For a given state *z* at time *t*, the multivariate high-gamma power signal *x* is modeled as

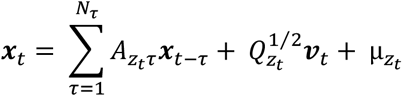

where μ*_z_* is a state-dependent bias. Probabilistic inference on observed neural data determines the unobserved latent parameters: the time series of network states *z_t_*, their transition probabilities 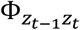
, and the linear dynamics parameters of each state {*A_z_, Q_z_*, μ*_z_*}.

The model was trained with iterative estimation of the state time series and state dynamics across all patients using the Baum-Welch expectation-maximization algorithm^71,72^. Initial conditions for *A* and *Q* were informed by the lagged correlation of multivariate AR clustering^73,74^. The state-dependent biases μ*_z_* were drawn randomly from a standard Gaussian. State transitions were sampled from a uniform prior.

### Training, Validation, and Testing

A random 80% of trials from each patient were used for training the latent dynamical parameters. 10-fold validation on the training set was used to select hyperparameters. The model contains 2 hyperparameters that constrain its architecture: model order *N*_τ_ and state number *N*_*s*_. Both were evaluated with log-likelihood and AIC. The return for increasing model order plateaued after *N*_τ_ *=* 3. Additional states exceeding *N*_*s*_ = 6 trivially split the background state during inter-trial periods. We trained a second model with interaction terms forced to zero – otherwise, the architecture was preserved. This mean-only model converged to a stable solution, but its performance was inferior to the complete model featuring interactions between regions.

The remaining 20% of trials from each patient were allocated to the testing pool to assess model fit. Performance of the model, measured with log-likelihood, was equivalent on training and testing sets. In addition, we iteratively held out each patient from training to ensure that the model was not overfit to individual-level effects. We report the state sequences generated by the model for all trials in the testing set.

### Quantifying Interactions with Partial Direct Coherence

The ARHMM classifies dynamical states by the network connectivity encoded in their AR coefficients. These dynamics can be captured by partial directed coherence (PDC) in the frequency domain^75^ – a measure of Granger causal information flow. For each state z, the pairwise PDC between regions *j* and *k* is defined as

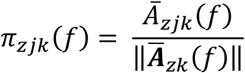

where

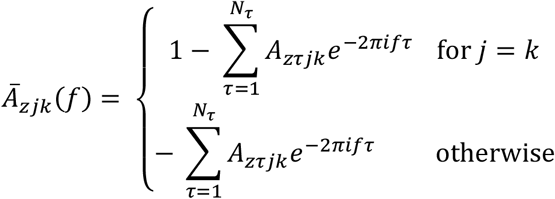

represents the transfer function at frequency *f*. In this manuscript, the directed graph for each state is shown with nodes representing regions and edges representing the broadband causal interactions.

## Acknowledgements

We thank all the patients who participated in this study; members of the Tandon lab (Patrick Rollo and Jessica Johnson); members of the Pitkow lab (Aram Giahi Saravani and Yicheng Fei); neurologists at the Texas Comprehensive Epilepsy Program (Jeremy Slater, Giridhar Kalamangalam, Omotola Hope, Melissa Thomas, Samden Lhatoo) who participated in the care of these patients; and all the nurses and technicians in the Epilepsy Monitoring Unit at Memorial Hermann Hospital who helped to make this research possible. This work was supported by the National Center for Deafness and Other Communication Disorders R01 DC014589 and F30 DC017083, National Institute of Neurological Disorders and Stroke U01 NS098981, National Science Foundation 1533664, National Science Foundation NeuroNex 1707400, National Science Foundation Career IOS-1552868, and the McNair Foundation.

---

**Supplemental Figure 1: SB-MEMA Movie**

Surface-based mixed-effects multilevel analysis of high-gamma power (112 patients, 13812 electrodes, 19465 trials). The model was calculated for 150ms wide windows in 10ms increments, time-locked separately to picture presentation and articulatory onset.

**Supplemental Figure 2:**
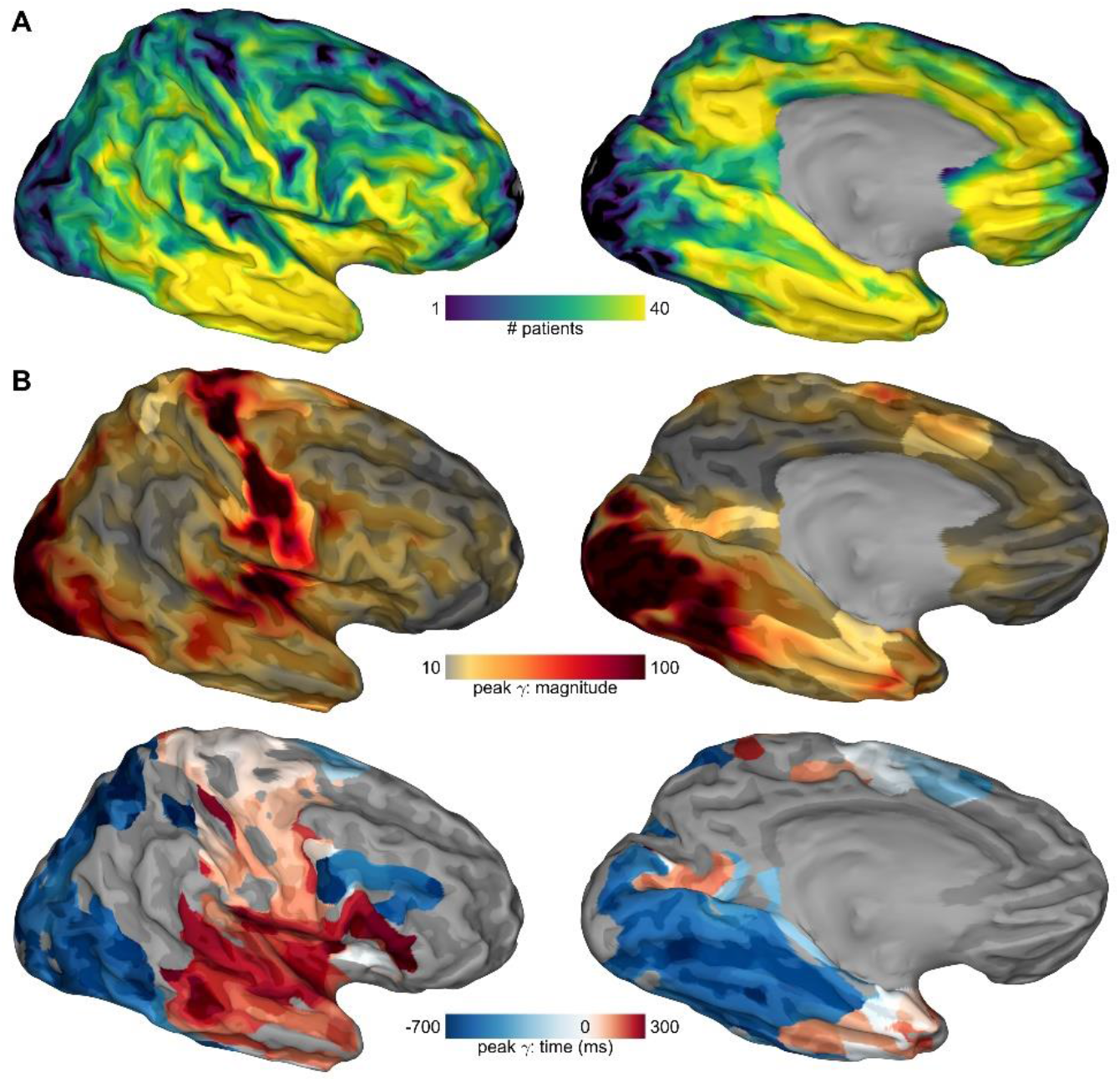
Spatiotemporal Atlas in the Language-Nondominant Hemisphere. Response dynamics in right language non-dominant hemisphere. **(A)** Aggregate cortical coverage in the patient population. **(B)** Surface-based mixed-effects multilevel analysis of high-gamma power.

**Supplemental Figure 5:**
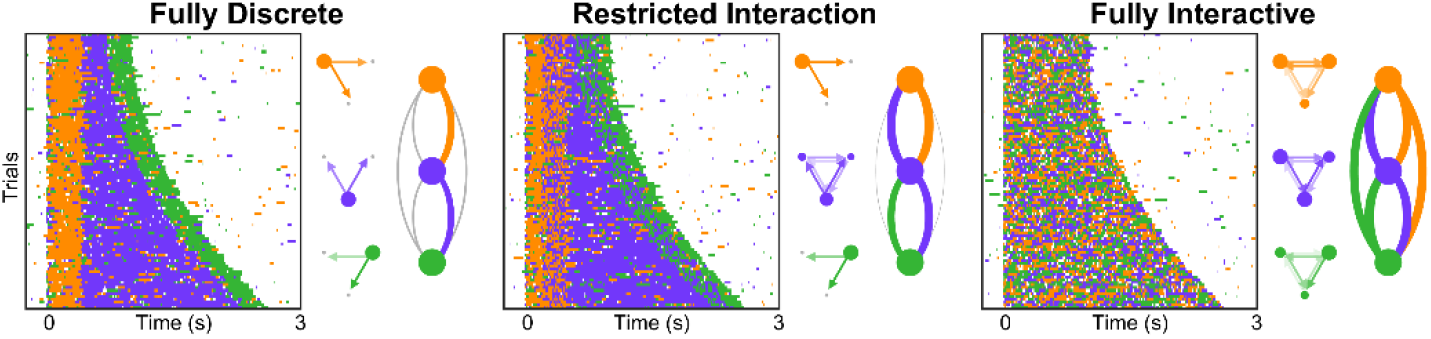
Linear Dynamical System Modeling of Discrete-Interactivity Axis. Theoretical models of speech production instantiated with switching linear dynamic systems. Each model features four unique states (white, orange, purple, and green) with pre-determined dynamics and state transition probabilities; these were used to generate 100 trials of data with varying reaction times at 3 regions. The state sequence raster, inter-regional interactions, and state transition probabilities are shown.

**Supplemental Figure 4:**
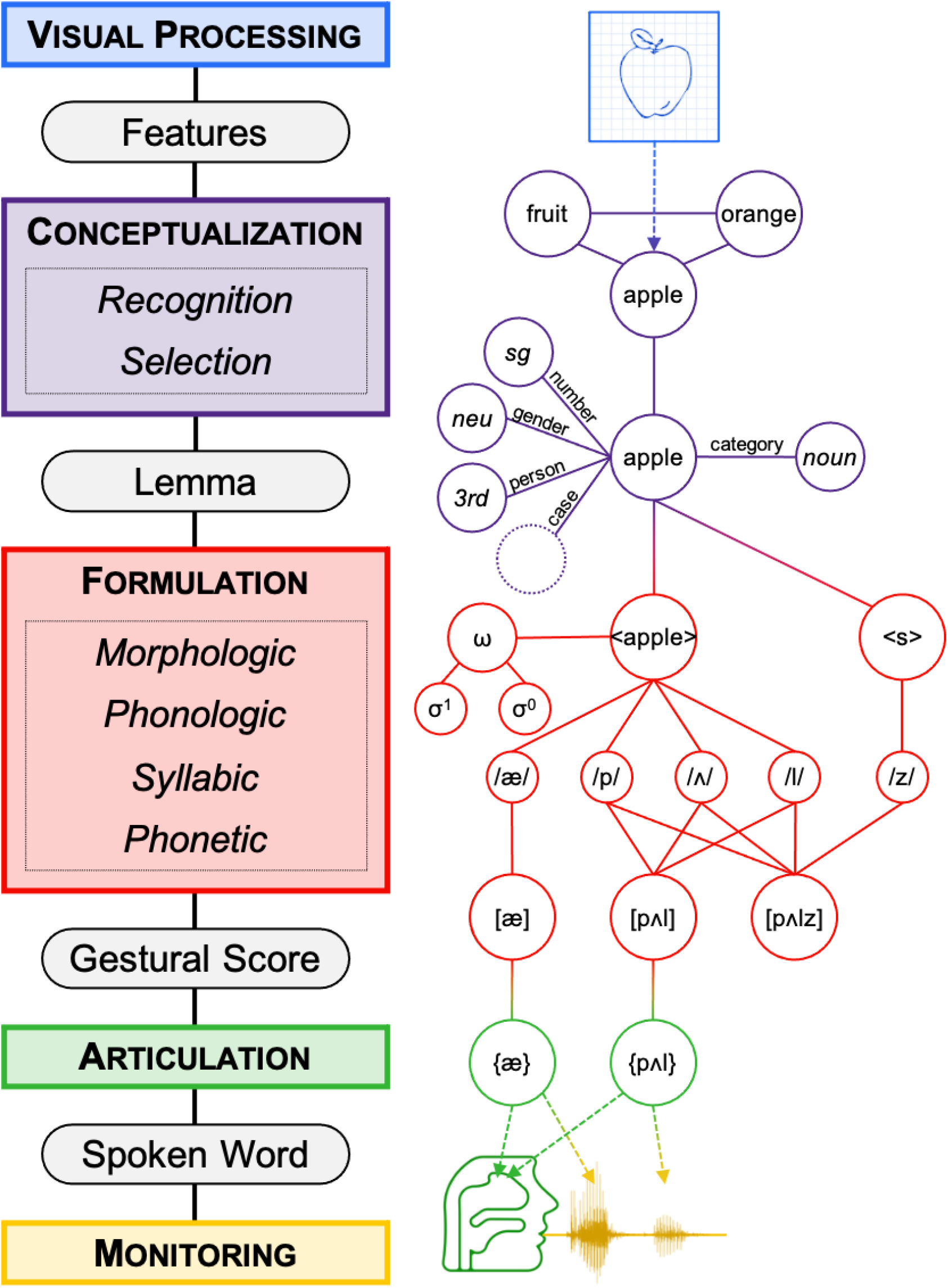
Indefrey & Levelt Model of Picture Naming. The five stages of cognitive processing during picture naming, as defined by Indefrey & Levelt. We have applied these stages to an example picture from the Snodgrass & Vanderwart stimuli.

**Supplemental Figure 5:**
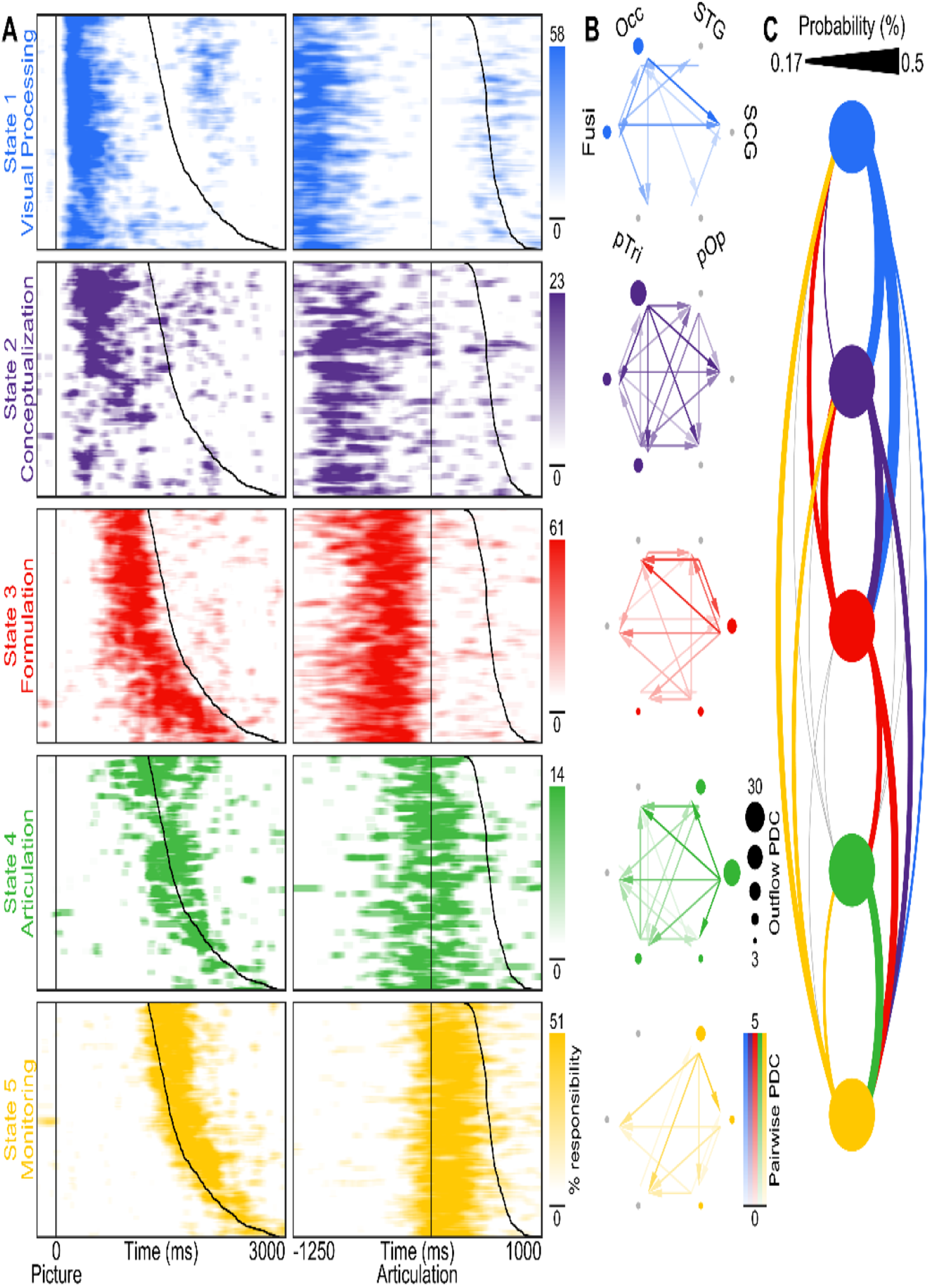
Nonlinear Dynamics during Scrambled Naming. Autoregressive hidden Markov model applied to single-trial high-gamma power during scrambled trials. **(A)** State sequences on held-out testing data. Each of the 5 active states is shown with a unique color. For each raster, all trials were sorted by reaction time in the left column and by articulatory duration in the right column. The maximum of each state color scale is set to 400% of the mean state density. **(B)** Interregional dynamics. The graph nodes represent distinct regions. The graph edges represent the pairwise partial directed coherence (PDC), a Granger causal measure of interaction derived from the unique dynamics that define each state. The size of each node is proportional to the total outflowing PDC. The significance of every pairwise interactional coefficient and each nodal outflow was evaluated via bootstrapping (n = 10000) at an alpha level of p < 0.001. No interactional coefficient in the residual state was significant. **(C)** State transition probabilities. The observed probability of transitioning “forwards” (e.g. blue to purple) is shown flowing downwards on the right; probability of transitioning “backwards” (e.g. yellow to green) is shown flowing upwards on the left. The lower threshold is set to the uniform probability of non-self-transitions.

**Supplemental Figure 6:**
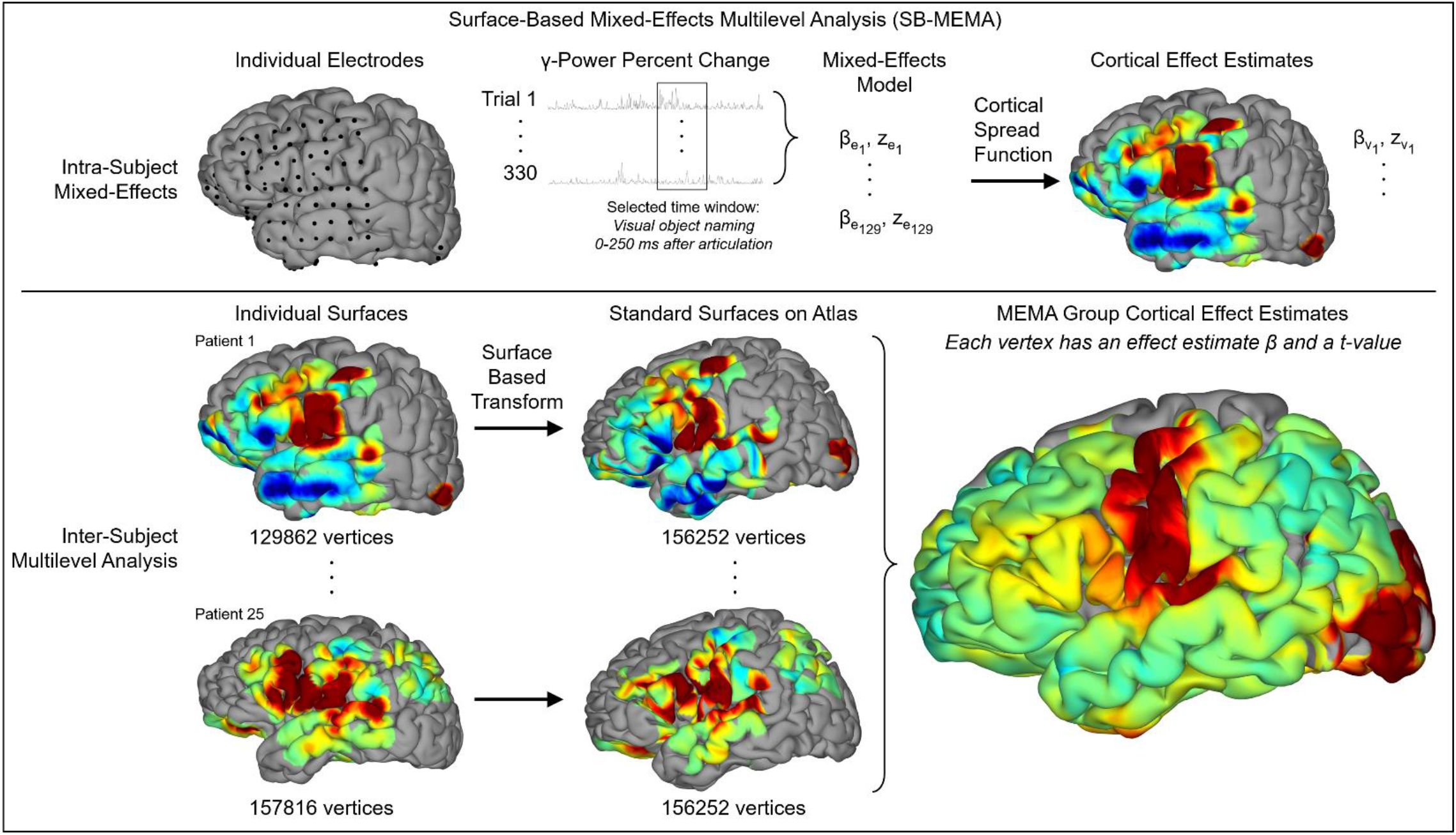
Surface-Based Mixed-Effects Multilevel Analysis (SB-MEMA) High-gamma power is extracted for all trials at each electrode and processed with a mixed-effects model. The effect size and confidence estimates are propagated onto the patient-specific pial surface via the cortical spread function. Patient surfaces are nonlinearly warped to the standard atlas using a surface-based transform. Data from all patients is integrated with a multilevel analysis in group space.

## Notes

### Competing Interest Statement

The authors have declared no competing interest.

## References

1. Indefrey, P. & Levelt, W. J. M. The spatial and temporal signatures of word production components. Cognition 92, 101–144 (2004).

2. Hillis, A. E. Aphasia: Progress in the last quarter of a century. Neurology (2007). doi:10.1212/01.wnl.0000265600.69385.6f

3. Warrington, E. K. The selective impairment of semantic memory. The Quarterly Journal of Experimental Psychology 27, 635–657 (1975).

4. Damasio, H.?;, Grabowski, T.J., Tranel, D.?;, Hichwa, R.D. & Damasio, A. R. A neural basis for lexical retrieval. Nature 380, 499–505 (1996).

5. Price, C. J. A review and synthesis of the first 20 years of PET and fMRI studies of heard speech, spoken language and reading. Neuroimage (2012). doi:10.1016/j.neuroimage.2012.04.062

6. Glasser, M. F. et al. A multi-modal parcellation of human cerebral cortex. Nature (2016). doi:10.1038/nature18933

7. Munding, D., Dubarry, A. S. & Alario, F. X. On the cortical dynamics of word production: a review of the MEG evidence. Language, Cognition and Neuroscience (2016). doi:10.1080/23273798.2015.1071857

8. Fedorenko, E. & Blank, I. A. Broca’s Area Is Not a Natural Kind. Trends in Cognitive Sciences (2020). doi:10.1016/j.tics.2020.01.001

9. Bassett, D. S. & Sporns, O. Network neuroscience. Nature Neuroscience (2017). doi:10.1038/nn.4502

10. Geschwind, N. Disconnexion syndromes in animals and man. Brain 88, 237 (1965).

11. Fedorenko, E. & Thompson-Schill, S. L. Reworking the language network. Trends in Cognitive Sciences 18, 120–126 (2014).

12. Crone, N. Functional mapping of human sensorimotor cortex with electrocorticographic spectral analysis. II. Event-related synchronization in the gamma band. Brain (1998). doi:10.1093/brain/121.12.2301

13. Bouchard, K. E., Mesgarani, N., Johnson, K. & Chang, E. F. Functional organization of human sensorimotor cortex for speech articulation. Nature 495, 327–332 (2013).

14. Sahin, N. T., Pinker, S., Cash, S. S., Schomer, D. & Halgren, E. Sequential processing of lexical, grammatical, and phonological information within Broca’s area. Science (80-.). 326, 445–9 (2009).

15. Gui, P. et al. Assessing the depth of language processing in patients with disorders of consciousness. Nat. Neurosci. (2020). doi:10.1038/s41593-020-0639-1

16. Rabinovich, M. I. Principles of brain dynamics: global state interactions. (MIT Press, 2012).

17. Kirst, C., Timme, M. & Battaglia, D. Dynamic information routing in complex networks. Nat. Commun. (2016). doi:10.1038/ncomms11061

18. Friston, K.J. Transients, metastability, and neuronal dynamics. Neuroimage (1997). doi:10.1006/nimg.1997.0259

19. Lowe, D. G. Distinctive image features from scale-invariant keypoints. Int. J. Comput. Vis. (2004). doi:10.1023/B:VISI.0000029664.99615.94

20. Snodgrass, J. G. & Vanderwart, M. A standardized set of 260 pictures: Norms for name agreement, image agreement, familiarity, and visual complexity. J. Exp. Psychol. Hum. Learn. Mem. 6, 174–215 (1980).

21. Brysbaert, M. & New, B. Moving beyond Kucera and Francis: a critical evaluation of current word frequency norms and the introduction of a new and improved word frequency measure for American English. Behav. Res. Methods 41, 977–90 (2009).

22. Vaden, K., Halpin, H. & Hickok, G. Irvine Phonotactic Online Dictionary, Version 2.0. (2009).

23. Rapp, B. & Goldrick, M. Discreteness and interactivity in spoken word production. Psychol. Rev. 107, 460–499 (2000).

24. Saravani, A. G., Forseth, K. J., Tandon, N. & Pitkow, X. Dynamic brain interactions during picture naming. eNeuro (2019). doi:10.1523/ENEURO.0472-18.2019

25. Roelofs, A. The WEAVER model of word-form encoding in speech production. Cognition 64, 249–284 (1997).

26. Dell, G. S., Schwartz, M. F., Martin, N., Saffran, E. M. & Gagnon, D. a. Lexical access in aphasic and nonaphasic speakers. Psychol. Rev. 104, 801–838 (1997).

27. Edwards, E. et al. Spatiotemporal imaging of cortical activation during verb generation and picture naming. Neuroimage 50, 291–301 (2010).

28. Conner, C. R., Chen, G., Pieters, T. A. & Tandon, N. Category specific spatial dissociations of parallel processes underlying visual naming. Cereb. Cortex 24, 2741–2750 (2014).

29. Kadipasaoglu, C. M. et al. Surface-based mixed effects multilevel analysis of grouped human electrocorticography. Neuroimage 101, 215–224 (2014).

30. Dubarry, A. S. et al. Estimating Parallel Processing in a Language Task Using Single-Trial Intracerebral Electroencephalography. Psychol. Sci. (2017). doi:10.1177/0956797616681296

31. Engel, A. K., Moll, C. K. E. E., Fried, I. & Ojemann, G. A. Invasive recordings from the human brain: clinical insights and beyond. Nat. Rev. Neurosci. 6, 35–47 (2005).

32. Jacobs, J. & Kahana, M. J. Direct brain recordings fuel advances in cognitive electrophysiology. Trends Cogn. Sci. 14, 162–171 (2010).

33. Llorens, A., Trébuchon, A., Liégeois-Chauvel, C. & Alario, F. X. Intra-cranial recordings of brain activity during language production. Frontiers in Psychology (2011). doi:10.3389/fpsyg.2011.00375

34. Halgren, E., Marinkovic, K. & Chauvel, P. Generators of the late cognitive potentials in auditory and visual oddball tasks. Electroencephalogr. Clin. Neurophysiol. (1998). doi:10.1016/S0013-4694(97)00119-3

35. Levelt, W. J. M., Roelofs, A. & Meyer, A. S. A theory of lexical access in speech production. Behavioral and Brain Sciences (1999). doi:10.1017/S0140525X99001776

36. Glaser, W. R. & Düngelhoff, F. J. The time course of picture-word interference. J. Exp. Psychol. Hum. Percept. Perform. (1984). doi:10.1037/0096-1523.10.5.640

37. Levelt, W. J. M. et al. The time course of lexical access in speech production: A study of picture naming. Psychol. Rev. (1991). doi:10.1037/0033-295X.98.1.122

38. Fromkin, V. a. The non-anomalous nature of anomalous utterances. Speech Errors As Linguist. Evid. 47, 27–52 (1973).

39. Garrett, M.. Levels of processing in sentence production. in Language production 177–200 (1980).

40. Dell, G. S. A spreading-activation theory of retreival in sentence production. Psychol. Rev. 93, 283–321 (1986).

41. Luria, A. R. Towards the mechanisms of naming disturbance. Neuropsychologia (1973). doi:10.1016/0028-3932(73)90028-6

42. Broca, P. Sur le siège de la faculté du langage articulé. Bull. la Société d’anthropologie Paris (1865). doi:10.3406/bmsap.1865.9495

43. Penfield, W. & Roberts, L. Speech and Brain Mechanisms. (Princeton University Press, 1959).

44. Hickok, G. Computational neuroanatomy of speech production. Nat. Rev. Neurosci. (2012). doi:10.1038/nrn3158

45. Binder, J. R. & Desai, R. H. The neurobiology of semantic memory. Trends Cogn. Sci. 15, 527–536 (2011).

46. Miozzo, M., Pulvermüller, F. & Hauk, O. Early parallel activation of semantics and phonology in picture naming: Evidence from a multiple linear regression MEG study. Cereb. Cortex (2015). doi:10.1093/cercor/bhu137

47. Conner, C. R., Kadipasaoglu, C. M., Shouval, H. Z., Hickok, G. & Tandon, N. Network dynamics of Broca’s area during word selection. PLoS One (2019). doi:10.1371/journal.pone.0225756

48. Kadipasaoglu, C. M. et al. Development of grouped icEEG for the study of cognitive processing. Front. Psychol. 6, 1–7 (2015).

49. Levelt, W. J. M. Models of word production. Trends Cogn. Sci. 3, 223–232 (1999).

50. Linderman, S., Nichols, A., Blei, D., Zimmer, M. & Paninski, L. Hierarchical recurrent state space models reveal discrete and continuous dynamics of neural activity in C. elegans. bioRxiv (2019). doi:10.1101/621540

51. Glaser, J. I., Whiteway, M., Cunningham, J. P., Paninski, L. & Linderman, S. W. Recurrent Switching Dynamical Systems Models for Multiple Interacting Neural Populations. bioRxiv (2020).

52. Hinton, G. E., McClelland, J. L. & Rumelhart, D. E. Distributed representations. Parallel Distributed Processing (1986).

53. Chen, G., Deng, Q., Szymczak, A., Laramee, R. S. & Zhang, E. Morse set classification and hierarchical refinement using conley index. IEEE Trans. Vis. Comput. Graph. (2012). doi:10.1109/TVCG.2011.107

54. Wada, J. & Rasmussen, T. Intracarotid Injection of Sodium Amytal for the Lateralization of Cerebral Speech Dominance. J. Neurosurg. 106, 1117–1133 (2007).

55. Tertel, K., Tandon, N. & Ellmore, T. M. Probing brain connectivity by combined analysis of diffusion MRI tractography and electrocorticography. Comput. Biol. Med. 41, 1092–1099 (2011).

56. Tandon, N. Cortical mapping by electrical stimulation of subdural electrodes: language areas. in Textbook of Epilepsy Surgery (ed. Luders, H.) 1001–1015 (McGraw Hill, 2008).

57. Oldfield, R. C. The assessment and analysis of handedness: The Edinburgh inventory. Neuropsychologia 9, 97–113 (1971).

58. Kaplan E, Goodglass H W.S. The Boston Naming Test. (Lea and Febiger, 1983).

59. Dale, A. M., Fischl, B. & Sereno, M. I. Cortical surface-based analysis. I. Segmentation and surface reconstruction. Neuroimage 9, 179–94 (1999).

60. Cox, R. W. AFNI: Software for analysis and visualization of functional magnetic resonance neuroimages. Comput Biomed Res 29, 162–173 (1996).

61. Pieters, T. A., Conner, C. R. & Tandon, N. Recursive grid partitioning on a cortical surface model: an optimized technique for the localization of implanted subdural electrodes. J. Neurosurg. 118, 1086–1097 (2013).

62. Forseth, K. J. et al. A lexical semantic hub for heteromodal naming in middle fusiform gyrus. Brain (2018). doi:10.1093/brain/awy120

63. Tandon, N. et al. Analysis of Morbidity and Outcomes Associated With Use of Subdural Grids vs Stereoelectroencephalography in Patients With Intractable Epilepsy. JAMA Neurol. (2019).

64. Gonzalez-Martinez, J. et al. Technique, Results, and Complications Related to Robot-Assisted Stereoelectroencephalography. Neurosurgery 78, 169–180 (2016).

65. Gonzalez-Martinez, J. et al. Stereotactic placement of depth electrodes in medically intractable epilepsy. J. Neurosurg. 120, 639–644 (2014).

66. Bruns, A., Eckhorn, R., Jokeit, H. & Ebner, A. Amplitude envelope correlation detects coupling among incoherent brain signals. Neuroreport 11, 1509–1514 (2000).

67. Saad, Z. S. & Reynolds, R. C. Suma. Neuroimage 62, 768–773 (2012).

68. Fischl, B., Sereno, M. I. & Dale, a M. Cortical Surface-Based Analysis II: Inflation, Flattening, and a Surface-Based Coordinate System. Neuroimage 9, 195–207 (1999).

69. Argall, B. D., Saad, Z. S. & Beauchamp, M. S. Simplified intersubject averaging on the cortical surface using SUMA. Hum. Brain Mapp. 27, 14–27 (2006).

70. Chen, G., Saad, Z. S., Nath, A. R., Beauchamp, M. S. & Cox, R. W. FMRI group analysis combining effect estimates and their variances. Neuroimage 60, 747–765 (2012).

71. Baum, L. E., Petrie, T., Soules, G. & Weiss, N. A Maximization Technique Occurring in the Statistical Analysis of Probabilistic Functions of Markov Chains. Ann. Math. Stat. (1970). doi:10.1214/aoms/1177697196

72. Dempster, A. P., Laird, N. M. & Rubin, D. B. Maximum Likelihood from Incomplete Data Via the EM Algorithm. J. R. Stat. Soc. Ser. B (1977). doi:10.1111/j.2517-6161.1977.tb01600.x

73. Morf, M., Vieira, A. & Kailath, T. Covariance Characterization by Partial Autocorrelation Matrices. Ann. Stat. (1978). doi:10.1214/aos/1176344208

74. Ding, M., Bressler, S. L., Yang, W. & Liang, H. Short-window spectral analysis of cortical event-related potentials by adaptive multivariate autoregressive modeling: Data preprocessing, model validation, and variability assessment. Biol. Cybern. (2000). doi:10.1007/s004229900137

75. Baccalá, L.A. & Sameshima, K. Partial directed coherence: A new concept in neural structure determination. Biol. Cybern. (2001). doi:10.1007/PL00007990

